## Supplemental Video Captions for "Open-source, high performance miniature multiphoton microscopy systems for freely behaving animals"

### Supplemental Videos

**Video 1: CA1 Recording**

This video demonstrates the recording presented in Figure 4. The FOV of the microscope is seen in the top-left image- excitatory neurons in hippocampal area CA1 are expressing CGaMP7f. Images were motion corrected in Suite2P and there is a ~ 1s rolling average applied to the signals, along with a ~ 0.5s maximum projection. Top-left shows the behavior of the animal, with a superimposed trace of the location, color coded by the animal speed. Individual ROIs were extracted from the calcium imaging data and those signals are plotted, synchronized to the free behavior of the animal. Lastly, the speed is quantified and plotted as a function of time as well.

**Video 2: RSC Recording**

This video shows the results from the 2-color cortical imaging of RSC presented in Figure 5. The animal is expressing GCaMP6f in L2/3 neurons, along with an activity-dependent mCherry cFos reporter. The strong brightness of the FOV shown in the top left is intentionally done for visualization purposes, as to demonstrate the ability of the microscope to resolve dynamics from dendrites. In reality, the somatic signals are well within the dynamic range of the detectors. Images are processed in the same way as Video 1. As in Video 1, the animal behavior is shown as well, time-synced with the UCLA 2P Miniscope recording, overlayed with the extracted position and motion. Activity from ROIs are shown, as well as the animal speed over the course of the experiment.

**Video 3: DG Recording**

This video is a detailed view of the deep recording of granule layer of dentate gyrus presented in Figure 6. The FOV expressing GCaMP8f is approximately 620 μm deep through an intact hippocampus. Processing is done in the same way as Videos 1 and 2 except an additional signal filter is applied to remove a minor artifact from the significantly weaker fluorescent signals. Animal behavior is shown, along with the activity traces from neurons as well as a quantification of the animal speed.
